# Systematic analysis of alternative splicing in time course data using Spycone

**DOI:** 10.1101/2022.04.28.489857

**Authors:** Chit Tong Lio, Zakaria Louadi, Amit Fenn, Jan Baumbach, Tim Kacprowski, Markus List, Olga Tsoy

## Abstract

During disease progression or organism development, alternative splicing (AS) may lead to isoform switches (IS) that demonstrate similar temporal patterns and reflect the AS co-regulation of such genes. Tools for dynamic process analysis usually neglect AS. Here we propose Spycone (https://github.com/yollct/spycone), a splicing-aware framework for time course data analysis. Spycone exploits a novel IS detection algorithm and offers downstream analysis such as network and gene set enrichment. We demonstrate the performance of Spycone using simulated and real-world data of SARS-CoV-2 infection.

## Background

Changes in alternative splicing (AS) lead to differential abundance of gene isoforms between experimental conditions or time points. If the relative abundance of two isoforms of a gene changes between two conditions or time points, this behavior is called isoform switching (IS). IS has a functional impact on the gene when the two switching isoforms perform different functions or when they have different interaction partners. Vitting-Seerup and Sandelin showed that IS changes the functions of 19% (N=2352) of genes with multiple isoforms in cancer, most of them leading to a protein domain loss [1]. In cardiovascular disease, the IS of Titin causes clinical symptoms of dilated cardiomyopathy [2,3]. Therefore, detection and functional interpretation of IS events is a promising strategy to reveal the mechanism of disease development.

However, the above examples refer to molecular snapshots of dynamic processes. In order to study such dynamic processes, like disease progression, we need time course data. By identifying groups of genes with similar temporal expression or AS/IS patterns, we can dissect the disease progression into mechanistic details. A study of mouse retinal development has shown that genes with similar temporal exon usage patterns shared similar biological functions and cell type specificity [4]. However, existing tools for AS analysis mostly focus on a single condition or two conditions from snapshot experiments. Tools developed for time course data analysis, e.g., TiCoNE [5], moanin [6], TimesVector-web [7]focus on gene expression level. TSIS, the only available tool to perform AS time course analysis, detects IS events whose effect lasts across several time points [8]. However, TSIS treats all IS events similarly, independent of their expression level. As a result, TSIS emphasizes isoforms with low expression while isoforms with comparably high expression levels are expected to be more involved in biological processes.

We introduce Spycone, a splicing-aware framework for systematic time course transcriptomics data analysis. It employs a novel IS detection method that priorizies isoform switches between highly expressed isoforms over those with minor expression levels, thus focusing on biologically relevant changes rather than transcriptional noise. Spycone operates on both gene and isoform level. For isoform-level data, the total isoform usage is quantified across time points. We have incorporated clustering methods for grouping genes and isoforms with similar time course patterns, as well as network and gene set enrichment methods for functional interpretation of clusters. The IS genes within the same clusters are expected to interact cooperatively with other functionally related genes. Thus, we hypothesize that disease mechanisms or developmental changes can be identified with network and functional enrichment methods. We compare the performance of Spycone and TSIS on a simulated and real-world dataset. On the latter we demonstrate how Spycone identifies network modules that are potentially affected by alternatively spliced genes during SARS-CoV-2 infection.

## Results

### Spycone overview

Spycone is available as a python package that provides systematic analysis of time course transcriptomics data. Figure 1 shows the workflow of Spycone. It uses gene or isoform expression and a biological network as an input. It employs the sum of changes of all isoforms relative abundance (total isoform usage) [9] (see Methods section), i.e. the sums of pairwise changes in relative isoform abundance, across time points to detect IS events. It further provides downstream analysis such as clustering by total isoform usage, gene set enrichment analysis, network enrichment, and splicing factors analysis. Visualization functions are provided for IS events, cluster prototypes, network modules, and gene set enrichment results.

**Figure 1.**
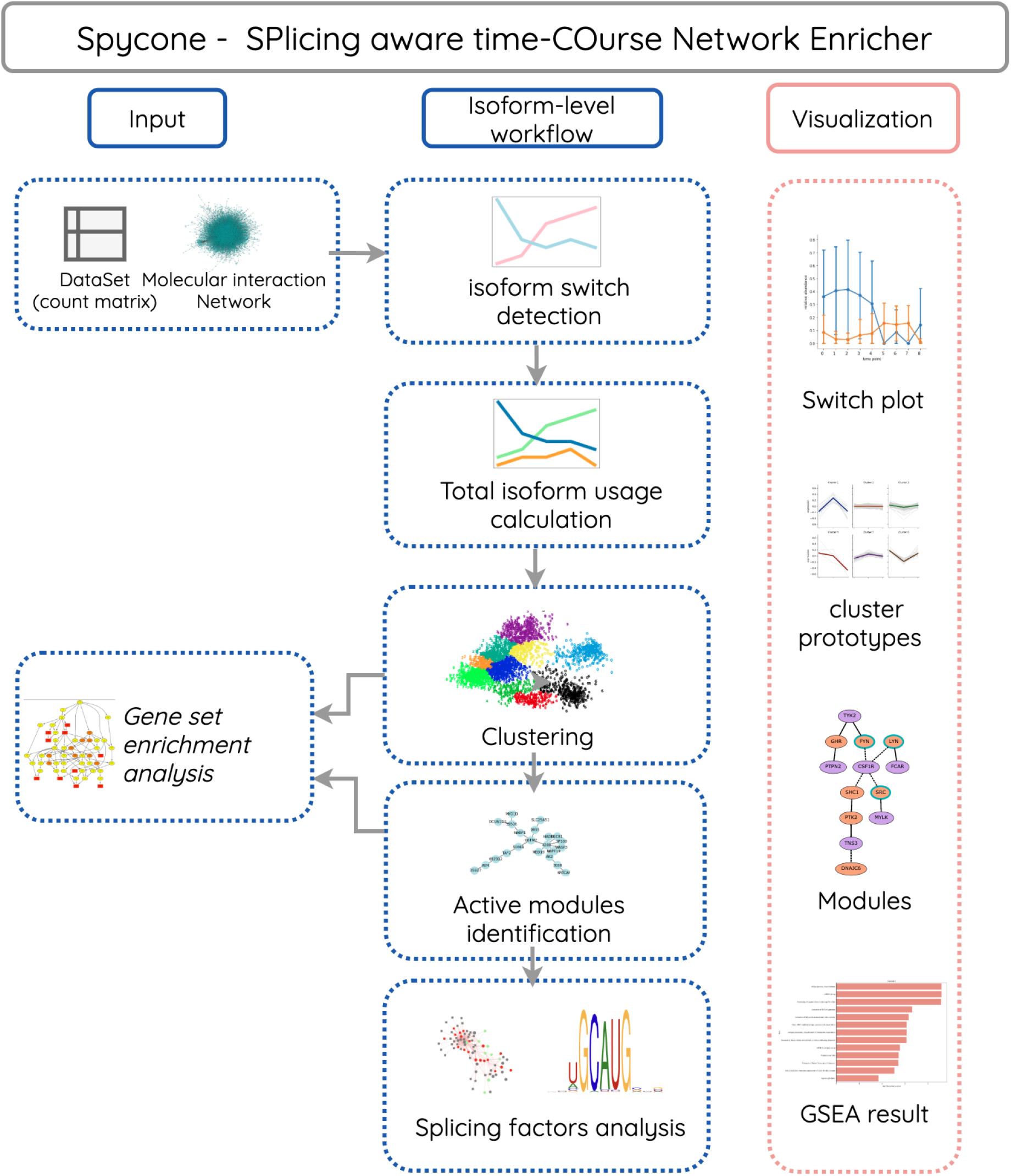
Overview of the Spycone architecture. Spycone takes count matrices and biological networks as input. We provide isoform-level functions such as isoform switch detection and total isoform usage calculation. Users could also cluster the gene count matrix directly. For downstream analysis, we integrated multiple clustering algorithms and an active modules identification algorithm (DOMINO). We also implemented splicing factors analysis for isoform-level data. Finally,visualizations are provided to better evaluate and interpret the results.

### IS detection

We propose novel metrics for the detection and selection of significant IS across time. IS events are described as a change of the isoform distribution between two conditions (time points). To detect an IS, our algorithm first searches for switch points, i.e. a specific time point where two isoform expression time courses intersect.

The main challenges to detect time course IS are: 1) most genes have multiple isoforms, the changes of the relative abundance can be due to factors other than AS, e.g. RNA degradation. 2) Most IS have multiple switch points, with different magnitudes of change in abundance; We need to consider how prominent the changes in abundance are to be recognized as an IS event. 3) Most genes have multiple lowly expressed isoforms that constitute noise and might not be biologically relevant. An ideal IS detection tool, therefore, should prioritize IS events according to their expression level (Fig S1).

Spycone overcomes these challenges by using a novel approach to detect IS events. Spycone uses two metrics: a p-value, and event importance. The p-value is calculated by performing a two-sided Mann-Whitney U-test between relative abundance before and after the switch point among the replicates. Event importance is the average of the ratio of the relative abundance of the two switching isoforms to the relative abundance of the isoform with the highest expression between the switching time points (see Methods section). Examples of high and low event importance are illustrated in Fig.2. The event importance will be highest when an IS includes the highest expressed isoform. Similarly, event importance will be low if an IS occurs between two lowly expressed isoforms. We also provide different metrics to comprehensively assess features of the IS events including switching probability, difference of abundance before and after switching and a dissimilarity coefficient (see Methods section).

**Figure 2.**
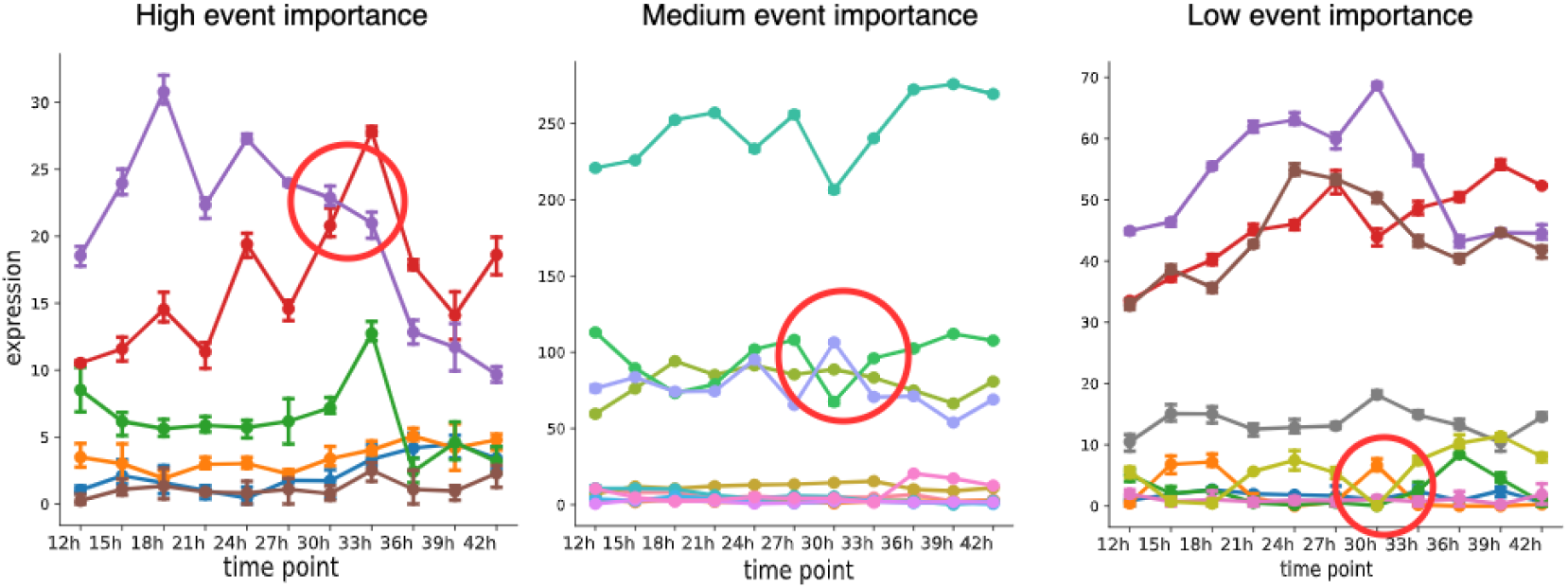
Plots showing the examples of three levels of event importance. Each plot contains all isoforms of a gene. The red circle indicates the IS events with the corresponding level of event importance.

### Clustering analysis for identifying co-spliced genes

Similarly to how transcription factors co-regulate sets of genes, in the context of AS, the splicing events of a subset of genes are co-regulated by splicing factors [10]. For genes with important IS events (identified as described above), we want to quantify the impact of splicing regulation between two time points. To this end, Spycone clusters genes by changes in total isoform usage over time to identify co-spliced genes. A previous study showed that clustering performance is highly dependent on the dataset and the clustering method [11]. Therefore, Spycone offers various clustering techniques, including agglomerative clustering (hierarchical clustering) [12], K-Means clustering [13], K-Medoids clustering [14], DBSCAN [15], OPTICS [16], and various distance metrics such as euclidean distance, Pearson distance, as well as tslearn [17] for calculating the dynamic time warping distance metric.

With temporal patterns of the clusters, Spycone dissects context-specific processes in terms of AS. In order to gain functional knowledge of the clusters, Spycone offers GSEApy [18] and NEASE [19] for gene set enrichment analysis. The former conducts classical enrichment analysis for multiple ontologies and pathway databases [20]. The latter combines information from protein-protein interaction (PPI) and domain-domain interaction networks, and allows to predict functional consequences of AS events caused by a set of IS genes.

### Active modules identification

Genes with consistent temporal patterns are thought to be functionally related in terms of co-regulation, molecular interactions, or participation in the same cellular processes. To uncover the underlying mechanism that is represented by a temporal pattern, Spycone projects the results of the clustering analysis on a molecular interaction network for active modules identification, i.e. for detection of subnetworks enriched in genes affected by IS. We incorporated DOMINO [21] as it has been previously demonstrated the best performance for this task [22]. To elucidate the functional impact of IS events, we further leveraged domain-domain interaction information from the 3did database [23]. Spycone identifies domains lost/gained during IS, which might indicate a functional switch, and affected edges in the PPI network. This provides additional insights about the functional consequences of time course IS.

### Splicing factor analysis

Spycone also provides splicing factor analysis using co-expression and RNA-binding protein motif search. Splicing factors are a group of RNA-binding proteins that regulate splicing of genes. We assume that the expression of splicing factors that are responsible for an IS event correlates with the relative abundance of participating isoforms. Spycone calculates the correlation between the expression value of a list of RNA-binding proteins derived from ENCODE eCLIP data [24,25] and the relative abundance of isoforms involved in IS. We implemented position-specific scoring matrices (PSSM) of RNA-binding protein motifs to calculate and detect the potential binding sites along the sequence of the targeted isoforms (see Method section).

### Evaluation using simulated data

To evaluate the performance of Spycone, we compared its performance (precision and recall) to TSIS using simulated data. TSIS provides an option to filter for IS events that involve only the highest abundance isoform - we refer to the result after filtering as major_TSIS. We aimed to investigate whether the performance of TSIS improves when applying this option.

We use a hidden markov model to simulate the switching state of the genes at each time point (see Methods section). We simulated two models (Fig. S2): Model 1 allows only major isoforms, i.e. those with the highest abundance per gene, to be involved in IS events across time points; Model 2 allows IS to occur between isoforms with relative abundance higher than 0.3. We used Model 2 to show that neither tool is biased towards events that involve only major isoforms.

For both tools, we varied their parameters (difference of relative abundance), to investigate how this affects their precision and recall. We also considered different levels of variance of gene expression, namely 1, 5 and 10, across replicates to mimic the noise (Fig. 3).

**Figure 3.**
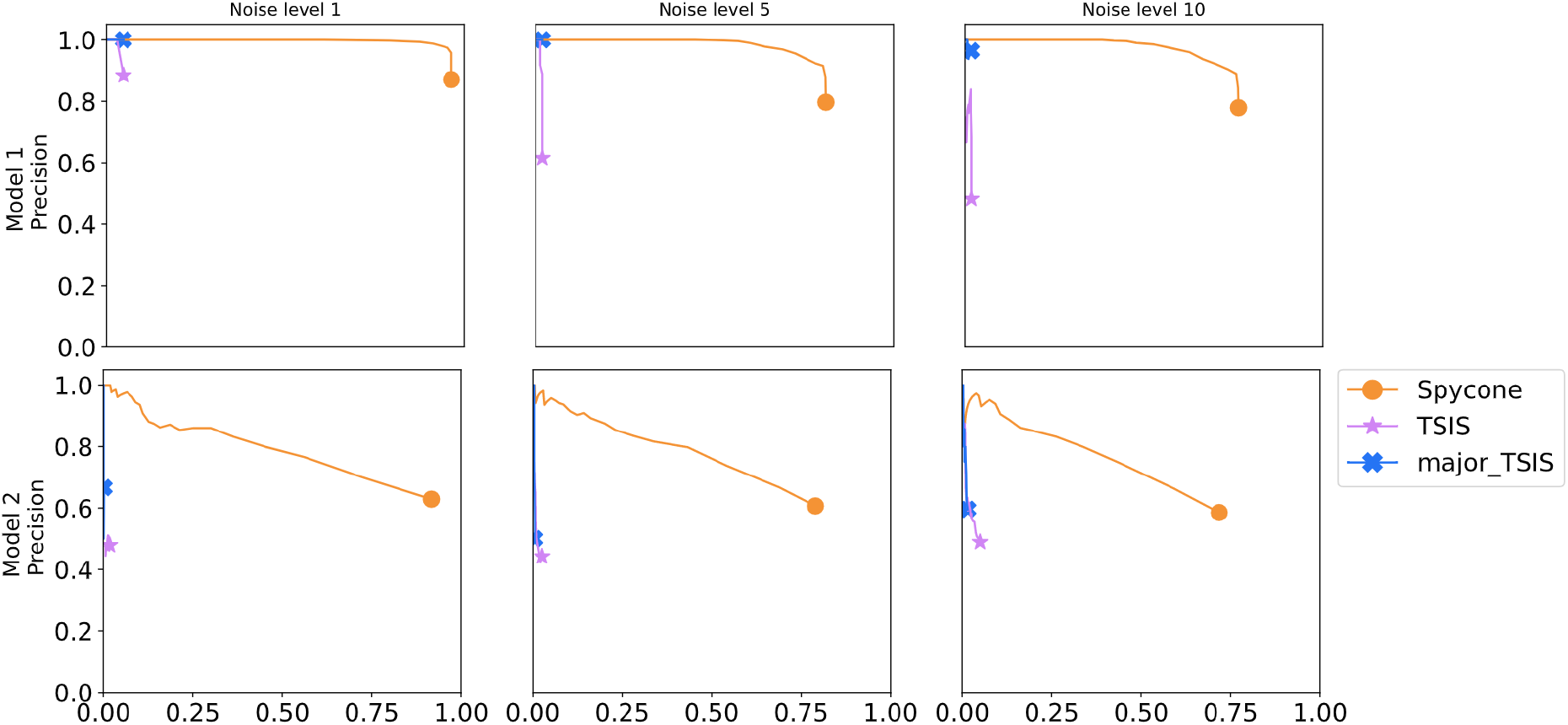
Precision and recall curves for Spycone, TSIS and major-isoforms filtered TSIS from simulated data of two models (rows) and three noise levels (columns).

In Model 1, Spycone achieved high precision and recall. The precision of TSIS dropped drastically with increasing recall. After filtering major events, TSIS’s recall reached 0.5. Spycone performs better in the setting with the highest noise level as it maintains high precision (0.95) and acceptable recall (0.75). In Model 2, Spycone achieved higher precision and recall than TSIS, however they dropped as the model became more stochastic. Moreover, TSIS has a higher algorithmic complexity of O(n*log(n)) than Spycone with a complexity of O(n), leading to a drastically lower runtime for Spycone in the range of a few minutes rather than hours (Fig. S3). In summary, Spycone outperforms TSIS in detecting IS events.

### Application to SARS-Cov2 infection data

We applied Spycone to an RNA-seq time course dataset of SARS-CoV-2 infected human lung cells [26]. The dataset contains 8 time points: 0, 1, 2, 4, 12, 16, 24 and 36 hours post-infection. We kept isoforms with transcripts per million (TPM) > 1 across all time points resulting in 36,062 isoforms for IS event detection with Spycone and TSIS. To call an IS significant, we used the following criteria: for Spycone, 1) switching probability > 0.5; 2) difference of relative abundance > 0.2 before and after the switch; 3) dissimilarity coefficient > 0.5; and 4) adjusted p-value < 0.05. For TSIS, we used 1) switching probability > 0.5; 2) difference of expression before and after switch > 10; 3) correlation coefficient < 0; 4) adjusted p-value < 0.05. The dissimilarity coefficient from Spycone and the correlation coefficient from TSIS are used to filter for IS events with negatively correlated isoforms. The values are chosen according to the performance on Model 2 simulated data with noise level 10 that showed the best precision. Spycone reported 915 IS events, of which 418 affected at least 1 protein domain. TSIS reported 985 events, of which 417 affected at least one protein domain. On gene level, Spycone reported 858 genes with IS events, TSIS reported 745 genes where 225 genes were found by both Spycone and TSIS (Fig. 4A).

**Figure 4.**
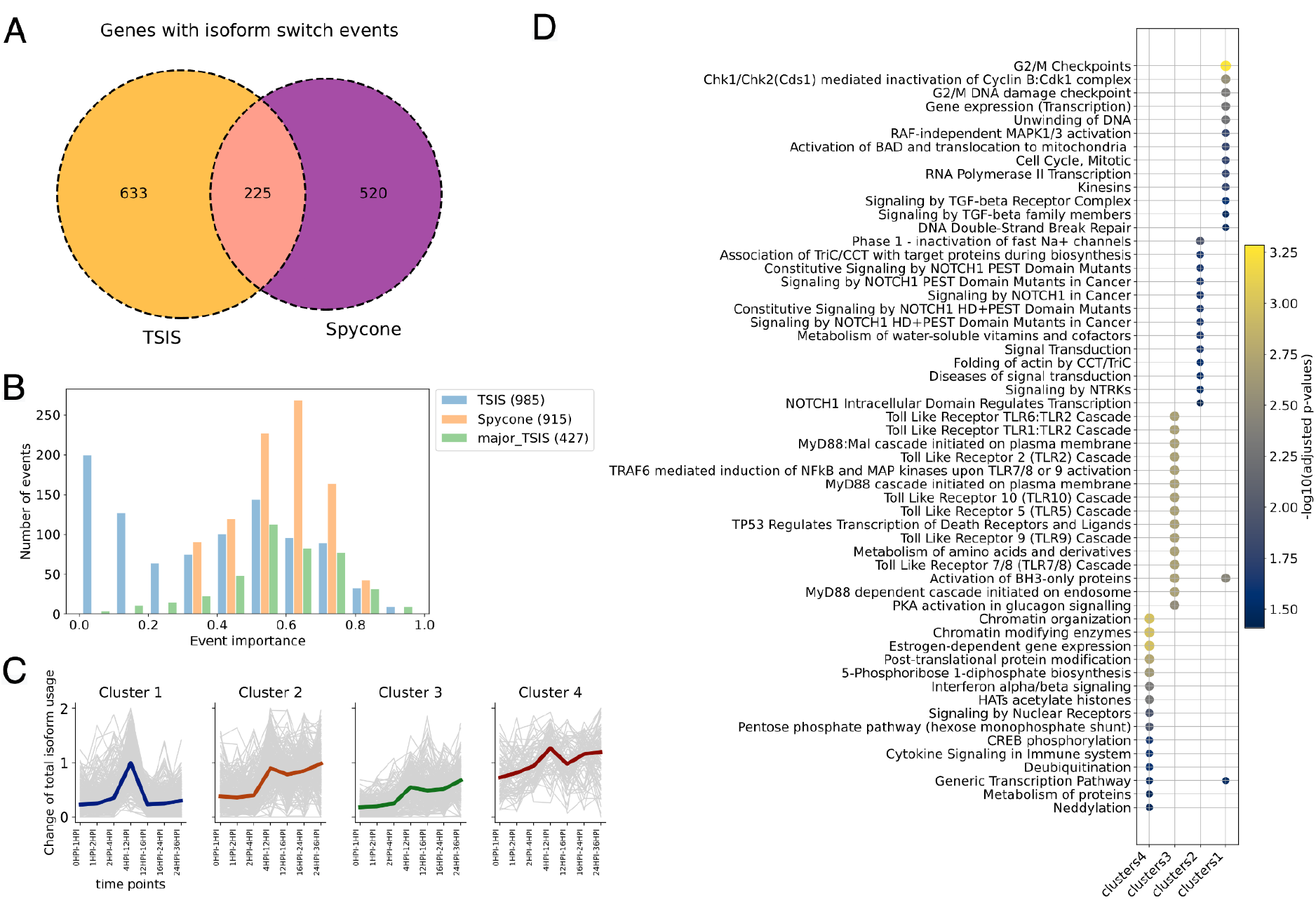
Comparing IS detection results from Spycone and TSIS A) Venn diagram showing the number of genes detected with isoform switch events by Spycone and TSIS (all). B) The distribution of the detected events in Spycone, TSIS and major_TSIS based on the event importance metric of Spycone. The number of events for each tool is indicated in brackets in the legend. C) Cluster prototypes (colored line) and all objects (gray lines) show the pattern of the change of total isoform usage across time points. D) NEASE enrichment results with Reactome pathways (y-axis) of the four clusters from Spycone (x-axis). The dotplot shows the −log10(adjusted p-value) from the hypergeometric test.

We then used the event importance metric to assess the ability of each method to detect IS events from higher abundance isoforms. We calculated event importance for IS events identified by Spycone, TSIS and major_TSIS (Fig. 4B). Spycone results include mostly events with high importance, while in TSIS events with low importance prevail. Table S1 shows the result for the SARS-CoV-2 dataset from the Spycone IS detection. Event importance has no clear prevalence towards overall gene expression and adjusted p-value. (Fig. S4, S5).

To exclude IS events with lowly expressed isoforms, we applied a filter of event importance higher than 0.3 to both Spycone and TSIS results. We calculated the change of total isoform usage of the IS genes across time points and employed Ward linkage hierarchical clustering. This led to four clusters with similar temporal patterns of changes in total isoform usage for Spycone (Figure 4C, Table S2) and four clusters for TSIS (Figure S4). Each cluster is represented by a cluster prototype, which is the median change of total isoform usage per pair of time points.

IS events that lead to domain gain or loss might break the interactions, hence rewiring the PPI network. Moreover, if the IS events belong to the same cluster, it indicates the synchronized gain or loss of interactions with particular pathways. Our goal is therefore to assess if IS events within clusters rewire interactions with particular pathways during SARS-CoV-2 infection. We performed AS-aware pathway enrichment analysis using NEASE with KEGG [27] and Reactome [28] pathway databases for results from Spycone (Fig. 4D, Table S3) and TSIS (Table S4, Fig S9). In addition, we performed classical gene set enrichment analysis using GSEApy. The results are not informative since only five terms are found in cluster 3 and zero in others.

Overall, clusters with similar prototypes from both tools are enriched in distinct pathway terms. For example, TSIS’s cluster 1 and Spycone’s cluster 1 have a strong peak between 4 and 12 hours post infection. Only TGF-beta signaling is commonly found in both tools. MAPK pathway and DNA damage checkpoint are enriched uniquely in Spycone. TSIS’s Cluster 2 and Spycone’s Cluster 3 have lower changes of total isoform usage overall. Spycone’s clusters showed more unique and relevant terms: 70 enriched Reactome terms in Spycone’s clusters and only 7 terms in TSIS’s clusters. TSIS’s cluster 3 and Spycone cluster 2 show an increase of change of total isoform usage after 12 hours post infection. Spycone’s cluster is enriched uniquely in protein folding chaperonin complex TriC/CCT and NOTCH signaling pathway. Finally, TSIS’s cluster 4 and Spycone’s cluster 4 have increasing changes of total isoform usage overall. TSIS’s cluster is enriched in mitosis-related pathways, cell cycle, and tubulin folding. Whereas in Spycone’s cluster 4 is found with signaling by PTK6, interferon, metabolism of proteins, pentose phosphate pathway etc.

Next, we detected active modules that show over-representation of IS genes from the same cluster based on DOMINO using a PPI network from BioGRID [29] (see Methods section). Detected active modules suggest the impact of splicing on regulatory cascades and cellular trafficking (Table 1).

**Table 1.** Related biological processes and pathways of the respective modules found in clusters.

| Cluster | Module | Related biological processes |
| --- | --- | --- |
| 1,<br>(Fig. 5A,<br>S7A) | 1 | RNA splicing and mRNA processing |
|  | 2 | cellular protein modification genes, signal transduction in the VEGF signaling pathway |
|  | 3 | positive regulation of protein ubiquitination |
|  | 4 | protein and intracellular trafficking genes |
|  | 5 | protein and intracellular trafficking genes |
| 2,<br>(Fig. 5B,<br>S7B) | 1 | transcription and mRNA splicing |
|  | 2 | Ras and Rho protein signaling transduction |
|  | 3 | ubiquitination |
|  | 4 | protein import into the nucleus |
|  | 5 | transition of cell cycle to G2/M phase |
| 3,<br>(Fig. 5C,<br>S7C) | 1 | regulation of transcription, cell cycle arrest and protein catabolic process |
|  | 2 | transmembrane receptor protein tyrosine kinase signaling pathway, in particular MAPK cascade and ERK cascade |
|  | 3 | protein ubiquitination |
|  | 4 | histone acetylation |
|  | 5 | organelle membrane fusion |
| 4,<br>(Fig. 5D,<br>S7D) | 1 | transcription and apoptotic processes |
|  | 2 | RNA splicing and processing |

### Splicing factor Analysis

Assuming that multiple IS events occurring between the same time points are co-regulated by the same splicing factor, we perform co-expression and motif analysis. The co-expression analysis yields thirteen significant RNA-binding proteins that are positively or negatively correlated with at least two isoforms of the same gene: in cluster 1 - FUBP3, HLTF, IGF2BP3, ILF3, RBFOX2, RBM22, SF3B1 and TAF15; in cluster 3 - IGF2BP3, RBM22, RPS6, SRSF7 and SUGP2; (|r| > 0.6 and adjusted p-value < 0.05) (Table S8).

To investigate whether the regulated exons, i.e. the lost or gained exons after IS events, show higher PSSM scores to a certain RNA-binding protein motif than the unregulated exons in a cluster, we applied motif enrichment analysis. We calculated PSSM scores along the flanking regions of the exons 5’ and 3’ boundaries and excluded the first and last exons in an isoform since these are often regulated by 5’-cap binding proteins and polyadenylation regulating proteins [30]. All exons in the switched isoforms within a cluster are categorized to 1) lost exons, 2) gained exons, and 3) unregulated exons for the analysis (Fig.6A, Table S9). RNA-binding proteins with multiple motifs are numbered with an underscore. Each motif is selected with a threshold where the false-positive rate is below 0.01. Position-specific log-odd scores higher than the corresponding threshold are obtained after calculating the PSSM scores of each motif for all exons (see Methods section). The ILF3_9 and ILF3_14 motifs show higher log-odd scores at the 5’ end of the lost/gain exons than of the unregulated exons in cluster 1 (one-sided Mann-Whitney U test p-value < 0.05) (Fig. 6A). HLTF_7 and SRSF7_1 motifs show higher log-odd scores at the 3’ end. (Fig. 6B)

## Discussion

AS regulates dynamic processes such as development and disease progression. However, AS analysis tools typically compare only two conditions and neglect how AS changes dynamically over time. Currently, the only existing tool for time course data analysis that accounts for splicing is TSIS. TSIS detects temporal IS events but is biased towards IS events between lowly expressed isoforms and does not offer features for downstream analysis which is important for interpreting the functional consequences of IS events.

Spycone, a framework for analysis of time course transcriptomics data, features a new approach for detecting temporal IS events and a new event importance metric to filter out lowly expressed isoforms. We demonstrate that Spycone’s IS detection method outperforms TSIS in terms of precision and recall based on simulated data. A key advantage of Spycone is that it explicitly considers how well IS events agree across replicates while TSIS considers averaged expression values among replicates and/or by natural spline-curves fitting. More specifically, Spycone uses a non-parametric Mann-Whitney U-test to test for significant IS and performs multiple testing correction to reduce type I error.

We have demonstrated the usability of Spycone by analyzing time course transcriptomics data for SARS-CoV-2 infection where we found affected signaling cascades. We performed NEASE enrichment on the clusters and compared the results from Spycone and TSIS. Spycone results are enriched in relevant terms such as MAPK pathway (cluster 1), NOTCH signaling (cluster 2), FGFRs and TLR pathways (cluster 3), and pentose phosphate pathway (cluster 4). NOTCH signaling pathways are found up-regulated in the lungs of infected macaques [31]. The MAPK pathway has a pro-inflammatory effect by interacting with SARS-CoV-2 downstream pathogenesis, especially in patients suffering from cardiovascular disease [32]. TLR 7/8 cascades are related to ssRNA, and there are studies supporting the association of TLR 7/8 with SARS-CoV-2 infection [33,34]. The pentose phosphate pathway is an alternative pathway of glycolysis that produces more NADPH oxidase. It is activated during SARS-CoV-2 infection in response to oxidative stress and the activation of immune response [35,36]. Spycone also detected the enrichment of pathways which association with SARS-CoV-2 infection has not been characterized yet: kinesins, signaling by NTRKs, degradation of AXIN, signaling by Hedgehog, and 5-phosphoribose 1-diphosphate biosynthesis.

The active modules extracted from the clusters highlight mechanisms involved in the host cell response to infection. In cluster 2, network enrichment analysis (module 1) (Figure 5B) revealed that interactions between three kinases (MAPK39, AURKC, and DCLK2) and a protein chaperone, HSP90AA1 is affected by IS. HSP90 is expressed under the ER stress caused by SARS-CoV-2 and its inhibitor is identified as a therapeutic inhibition target [37]. A previous study found that knock down of MAP3K9 reduced SARS-CoV-2 virus replication [38]. DCLK2 is differentially expressed in SARS-CoV-2 patients [39]. AURKC would be an interesting candidate to investigate for its role in SARS-CoV-2 infection.

**Figure 5.**
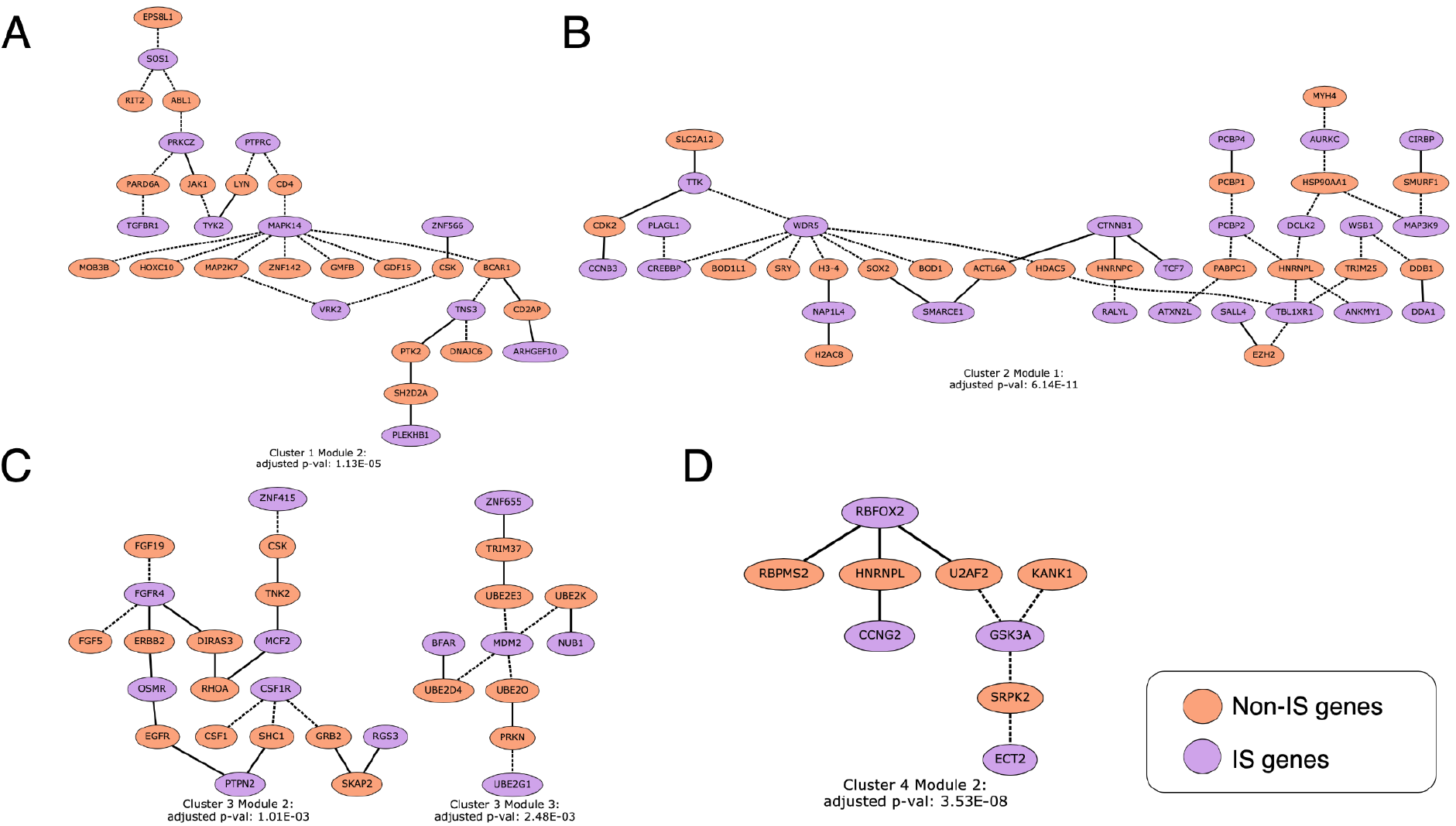
Spycone results in modules of the PPI network and their corresponding gene set enrichment results. Active network modules are identified using DOMINO. Each node represents a domain of a gene. Purple nodes are the isoform switched genes and orange nodes are non-IS genes from the PPI. Dashed edges are the affected interactions between the genes due to the loss/gain of domains during the IS events.

**Figure 6.**
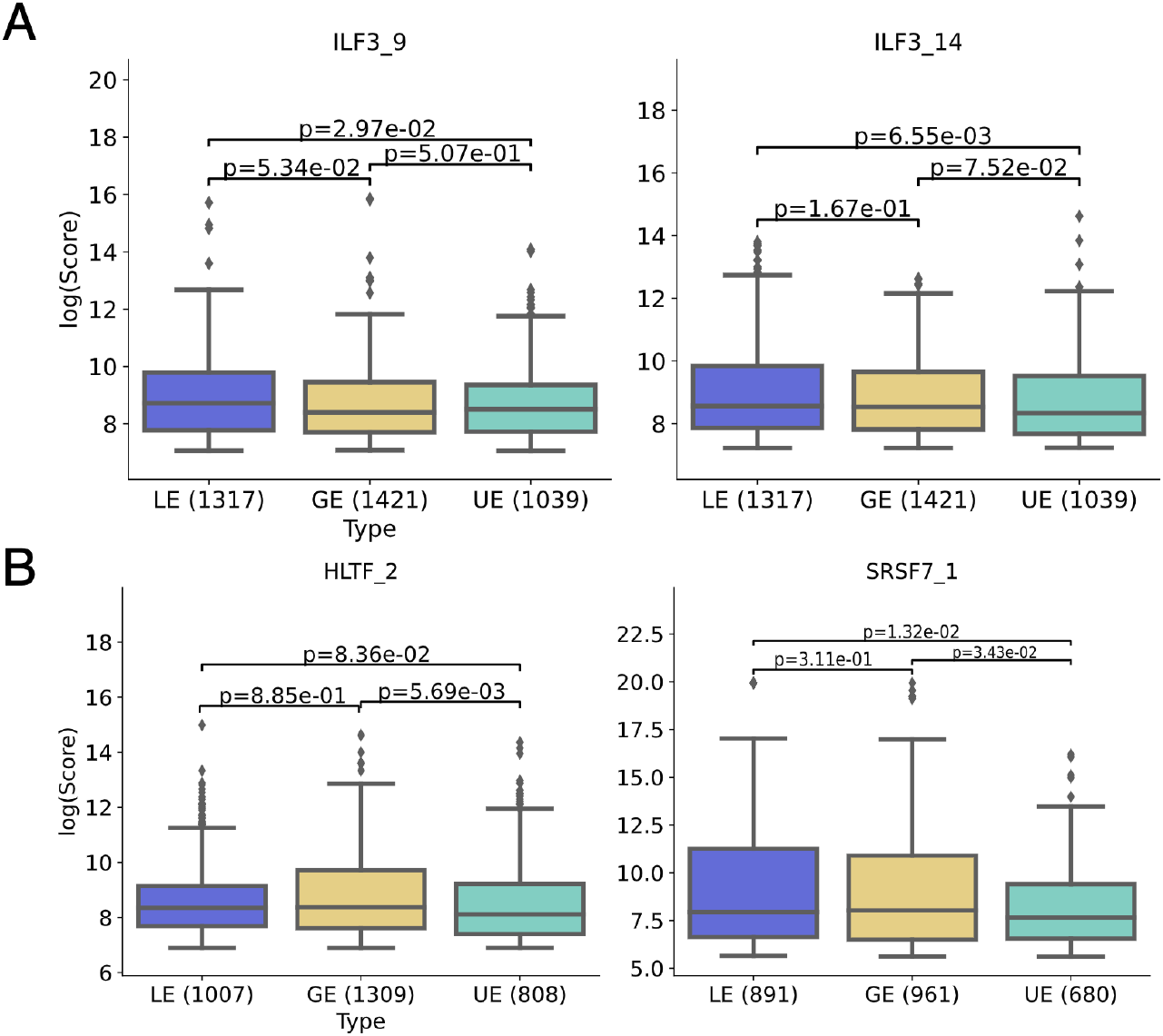
A) Boxplots showing the PSSM score difference between lost/gained exons and unregulated exons at the exon 5’ boundaries in logarithmic scale (one-sided Mann Whitney U test p-value < 0.05). B) Boxplots showing the PSSM scores difference between lost/gained exons and unregulated exons at the exon 3’ boundaries in logarithmic scale (one-sided Mann Whitney U test p-value < 0.05). Abbreviations: LE: Lost Exons; GE: Gained Exons; UE: Unregulated Exons.

Besides these three kinases, network enrichment analysis highlighted the general importance of kinases in infection development. e.g., JAK1, LYN, TYK2, PRKCZ (Figure 5). JAK1 is responsible for interferon signaling [40]. Inhibition of LYN reduces efficiency of SARS-CoV-2 virus replication [41]. TYK2, which is a key player for IFN signaling, has been associated with cytokine storms in SARS-CoV-2 patients [42]. IS events of kinases might cause major rewiring of the transduction cascade, which could lead to altered immune response, cell cycle control and promote viral replication.

Our analysis also suggests an important role of growth factor receptors (FGFR, EGFR, VEGF) and their downstream kinases. They are essential for viral infection since they modulate cellular processes like migration, adhesion, differentiation and survival. One example is that activation of EGFR in SARS-CoV-2 can suppress the IFN response and aid viral replication [43].

Another key finding is that E3 ubiquitin ligases are affected by IS. They are known to mediate host immune response by removing virus particles. Various virus species hijack the host E3 ubiquitin ligases in favor of viral protein production [44]. They are also involved in maintaining TMPRSS2 stabilization during virus entry to the host cells [45].

In splicing factor analysis, ILF3 and SRSF7 are identified as a splicing factor affecting the splicing of exons. ILF3 plays a role in antiviral response by inducing the expression of interferon-stimulated genes [46]. In another computational analysis, SRSF7 is also predicted to have binding potential with SARS-CoV-2 RNA [47].

Lastly, in order to get confident time course analysis results, one will need high resolution data in terms of number of time points and sample replicates. Consequently, at least three time points and three replicates are recommended in Spycone analysis. However, this criteria is rather met due to technical and economical restraints. Thus, Spycone also provides an option for a permutation test with only one replicate for the dataset under investigation. We demonstrated this usage in a tumor development dataset with one replicate (see supplementary information).

### Limitations

Spycone achieves high precision and considerably higher recall than the only competing tool TSIS. Nevertheless, the moderate recall we observe in particular in the presence of noise shows that there is further room for method improvement. In our simulation Model 2, where we allowed for isoform switches between minor isoforms, we observed a reduction in both precision and recall. Spycone identifies only two isoforms that switch per event, but in reality, an event could involve more than two isoforms. In the future, we should consider multiple-isoforms switches to handle more complex scenarios.

Spycone uniquely offers features for detailed downstream analysis and allows for detecting the rewiring of network modules in a time course as a result of coordinated domain gain/loss. This type of analysis is limited by the availability of the structural annotation. However, the current developments in computational structural biology that could expand the information about domains and domain-domain interactions e.g., AlphaFold2 [48], will greatly strengthen our tool. Lastly, our PSSM-based approach for splicing factor analysis does not allow us to investigate splicing factors that bind indirectly through other adaptor proteins, requiring further experiments that establish binding sites for such proteins.

Spycone was thus far applied exclusively to bulk RNA-seq data. When considering tissue samples, IS switches between time points could also be attributed to changes in cellular composition. An attractive future prospect is thus to apply Spycone for studying IS in single-cell RNA-seq data where dynamic IS events could be traced across cellular differentiation using the concept of pseudotime. However, the current single-cell RNA-seq technologies are limited in their ability to discern isoforms [49].

### Conclusion

With declining costs in next-generation sequencing, time course RNA-seq experiments are growing in popularity. Although AS is an important and dynamic mechanism it is currently rarely studied in a time course manner due to the lack of suitable tools. Spycone closes this gap by offering robust and comprehensive analysis of time course IS. Going beyond individual IS events, Spycone clusters genes with similar IS behavior in time course data and offers insights into the functional interpretation as well as putative mechanisms and co-regulation. The latter is achieved by RNA-binding protein motif analysis and highlights splice factors that could serve as potential drug targets for diseases. Using simulated and real data, we showed that Spycone has better precision and recall than its only competitor, TSIS, and that Spycone is able to identify disease-related pathways in the real-world data, as we demonstrated for SARS-CoV-2 infection. In summary, Spycone brings mechanistic insights about the role of temporal changes in AS and thus perfectly complements RNA-seq time course analysis.

## Materials and Methods

### Data Preprocessing

We demonstrated the performance of Spycone on RNA-seq data from SARS-CoV-2 infected human lung cells (Calu-3) with 8 time points and 4 replicates for each time point [26]. For the SARS-CoV-2 dataset, we used Trimmomatic v0.39 [50] to remove Illumina adapter sequences and low quality bases (Phred score < 30) followed by Salmon v1.5.1 [51] for isoform quantification with a mapping-based model, the human genome version 38, and an Ensembl genome annotation version 104.

### Protein-protein interaction network and Domain-domain interaction

A PPI network is obtained from BioGRID (v.4.4.208) [29] and a domain-domain interaction network from 3did (v2019_01) [23]. The edges of the PPI network are weighted according to the number of interactions found between the domains of the protein (nodes of PPI), given by the domain-domain interaction. Weighting PPIs with domain-based information can result in a functionally more interpretable network in diseases and pathways [52].

### Simulation

We used the SARS-CoV-2 dataset described above as a reference for setting the parameters of a negative binomial distribution of gene expression counts, as well as the parameters of Poisson distribution of the number of isoforms for each gene.

### First-order Markov chain

A first-order Markov chain is used for simulation of the gene states at each time point. In the simplest form, we defined two gene states: switched or unswitched. Change of the states along the time course depends on the transition probabilities.

We used a Dirichlet distribution to simulate relative abundance for each isoform of a gene. The relative abundance of an isoform is the ratio of the isoform expression to the total gene expression. The outcome of the Dirichlet distribution is *k*-dimensional vectors *x* with real numbers between 0 and 1 such that the sum of the elements in *x* is 1. This is suitable to simulate probability distribution of *k* categories. The Dirichlet distribution is defined as:

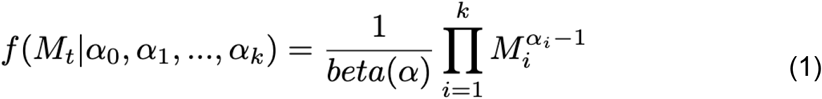

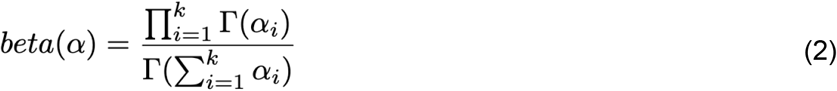

where the parameter α is a k-dimensional vector governing the distribution of the probabilities. In our case *k* equals the number of isoforms in a gene, where each isoform will be assigned an α_i_ value. The higher α_i_, the higher the probability of the isoform will be assigned an *a_i_* value.The higher *a_i_* the higher the probability of the isoform i.

In Model 1, where we assumed that switching isoforms are highly expressed, the α for switching isoforms are α = {1,2,…, *s*] * 10, s=*number of switching isoforms*, while α for the remaining isoforms are 1. To introduce switching events, the isoform probabilities of two highly expressed isoforms are swapped. For instance, if the isoform probabilities of the unswitched state for gene g with 5 isoforms are {0.03, 0.07, 0.1, **0.3, 0.5**}. Then the isoform probabilities for the switched state is {0.03, 0.07, 0.1, **0.5, 0.3**}.

In Model 2, where we assumed that isoforms with abundance higher than 0.3 have equal chances to switch, the α vector is α = {1,2,…,*k*} * 10, k=*number of all isoforms*,. To introduce switching events, the probabilities of two random isoforms will be swapped.

After we simulated abundances for each isoform, we multiplied it to a gene expression mean selected based on real life dataset to obtain the transcript expression mean (μ_i_). The gene expression means are randomly picked among the genes with the same number of isoforms from the real-world dataset.

We simulated time course data with 10 time points and 3 replicates using 10,000 genes. The transcript expression of replicates are sampled from normal distribution with a given transcript mean (μ_i_), and the variance is sampled from a gamma distribution as the following:

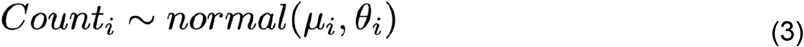

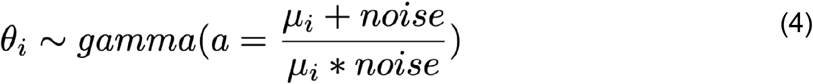

In order to simulate the differences generated for each individual experiment in real life, we tested on noise levels 1, 5 and 10. This setting can ensure isoforms with higher abundance will have a higher variance compared to those with lower abundance.

The data was analyzed with Spycone’s *detect_isoform_switch* function and TSIS’s *iso.switch* function. TSIS was further tested in two modes - TSIS and major_TSIS - major_TSIS uses the *max.ratio = TRUE* parameter.

### Detection of isoform switch events

#### Spycone

The first step of IS detection is to filter out transcripts that have an average TPM < 1 over all time points. Spycone then detects IS events based on relative abundance of the isoforms. The IS events are defined with the following metrics:

1. Switching points Switch points refer to the points where two time courses intersect in at least 60% of the replicates. For every pair of isoforms in a gene, Spycone detects all possible switch points for further analysis. For a dataset that has only one replicate, Spycone checks the intersection between isoform pairs in one replicate.
2. Switching probability As TSIS, Spycone calculates a switching probability for each IS event. A switching probability is the frequency of samples where isoform i is higher than isoform j before switch, and vice versa, the frequency of samples where isoform i is lower than of isoform j after switch. Time points between switching points are taken into account. If two isoforms switched at *t_x_* and *t_y_* the switching probability between isoform i and isoform j between time *t_x_* and *t_y_* is:

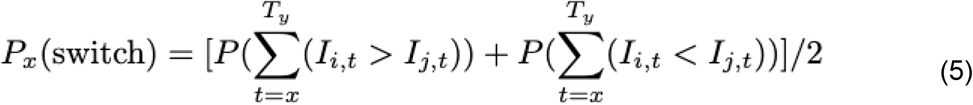
3. Significance of switch points If replicates are available, Spycone calculates the significance of a switch point by performing a two-sided Mann-Whitney U-test between relative abundance before and after the switch point similar to TSIS. For a dataset that has only one replicate, a permutation test is performed, where the time points within a time course are permuted. An empirical p-value is calculated to indicate the probability for the two switching isoforms to have a higher dissimilarity coefficient and higher difference of relative abundance before and after switch. Since the goal here is to select genes that have significant IS, Spycone takes the best switch point for further analysis with the smallest p-value. Other significant switch points will be reported as part of the result for users to investigate.
4. Difference of relative abundance To quantify the magnitude of changes during IS, Spycone calculates the average difference of relative abundance before and after a switch point. If replicates are available, Spycone calculates the average change of relative abundance. We selected a cutoff of 0.1, where the changes of the relative abundance accounts for at least 10% of the total gene expression. Difference of relative abundance between switching isoforms i and j is defined as:

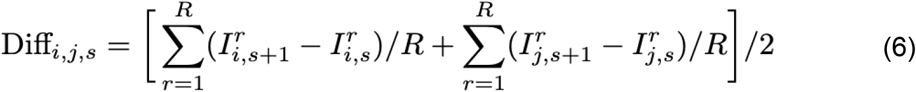

where s is a switch point; R is the number of replicates.
5. Event importance Event importance is a novel metric that accounts for the expression level of switching isoforms. We defined event importance as:

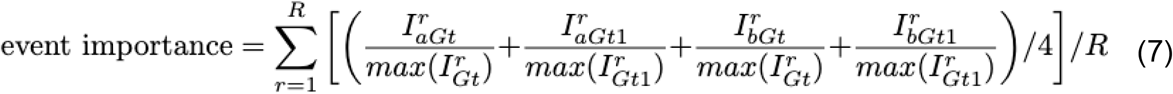

where 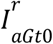 is the expression of isoform a of a gene *G* at time point *t0*; and *R* is the total number of replicates. Each I is normalized to the highest relative abundance 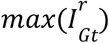 at the corresponding time point. For the analysis we used IS events with differences of relative abundance higher than 0.1 and event importance higher than 0.2.
6. Dissimilarity coefficient Dissimilarity coefficients d_i,j assess the dissimilarity of the time course between isoforms. It is calculated based on Pearson correlation r_i,j between time course i and j:

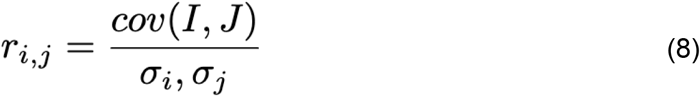

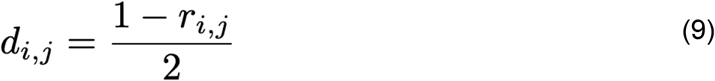 The higher coefficient, the less similar are the time courses.
7. Domain inclusion or exclusion We used the Pfam database v.35.0 [53] to map domains to isoforms. Spycone compares isoforms in the IS event with each other to define if there is a loss/gain of domain.
8. Multiple testing correction Finally, we implemented multiple testing corrections for IS detection. Available corrections are Bonferroni, Holm-Bonferroni and Benjamini-Hochberg false discovery rate (FDR). We use the Benjamini-Hochberg method as default.

### TSIS

To detect IS in TSIS, we used the following parameters: 1) the switching probability > 0.5; 2) difference before and after switch > 10; 3) interval lasting before and after at minimum 1 time point; 4) p-value < 0.05 and 5) Pearson correlation < 0. More detailed descriptions of parameters are found in [8]. The above parameters are set with defaults suggested by TSIS, except parameter 3), since we have a larger interval between time points (12 hours at maximum).

The computational method of TSIS utilizes 4 metrics [8]:

1. Switching probability It calculates the probability that one isoform is more or less abundant than the other isoform.
2. Difference of isoform expression It calculates the sum of the average differences of expression before and after the switch points.
3. Significance of switch points It calculates p-value from paired t-tests of the difference before and after switch points. Since TSIS does not correct for multiple testing. We further applied the p-values with the Benjamini-Hochberg method to obtain adjusted p-values.
4. Correlation coefficient Pearson correlation coefficient across all time points between two isoforms time course.

### Change of total isoform usage

Isoform usage measures the relative abundance of an isoform. Isoform usage of all isoforms from one gene are summed up to obtain the total isoform usage. We defined the change of total isoform usage as between two consecutive time points:

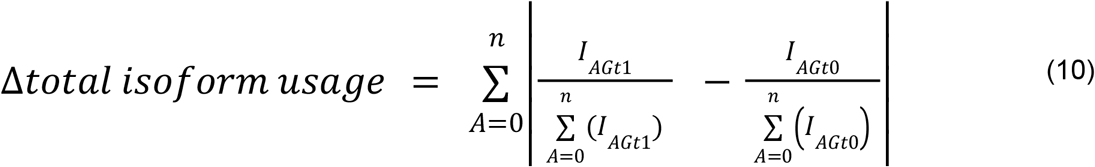

where *I* is the expression of isoform *A* of gene *G* at time points *t1 at t0;* and *n* is the total number of all isoforms for a gene *G*.

### Clustering analysis

The clustering algorithms are implemented using the *scikit-learn* machine learning package in python (v0.23.2) [54] and *tslearn* (v0.5.1.0) time course machine learning package in python [17]. The available algorithms are K-means, K-medoids, agglomerative clustering, DBSCAN, and OPTICS.

The number of clusters is chosen manually by visually checking the Ward distance dendrogram (Supplementary S6, 7).

### Gene set enrichment and network analysis

For enrichment analysis Spycone uses Gseapy and NEASE. Gseapy is a python wrapper of GSEA and Enrichr [20]. Gene set enrichment method is performed using Fisher’s exact test. NEASE [19]is an enrichment method for co-regulated alternative exons. We used NEASE with KEGG and Reactome pathways. For a seamless analysis, the newest version of the NEASE’s Python package (v1.1.9) is integrated with Spycone.

Spycone employs DOMINO (0.1.0) [21] for active module identification in PPI networks using default parameters.

### Splicing factor co-expression and motif enrichment analysis

List of splicing factors and their PSSMs are obtained from the mCross database (downloaded in 2022), currently only available for *Homo sapiens*) [25]. First, we filtered splicing factors with TPM > 1 in all time points. Next, we calculated the correlation between the relative abundance of each isoform and the expression of splicing factors. We filtered the pairs with correlation > 0.7 or < −0.7 and adjusted p-value < 0.05.

Finally, we performed motif enrichment analysis using the *motifs* module from the Biopython library [55]. The *motifs* module computes log-odd probability of a specific region in the genome to match the binding motif using the PSSM [56]. Hence, the higher the log-odd score, the more likely the binding. We compared these scores obtained from the lost, gained, and unregulated exons from the same clusters. A Mann-Whitney U test is performed on the sets of scores. Each motif threshold is selected using the *distribution* of the PSSM score over the frequency of nucleotides (background). The threshold is set at a false-positive rate < 0.01, meaning the probability of finding the motif in the background is less than 0.01.

## Supporting information

Supplementary information - use case 2

## Data Availability

The SARS-CoV-2 infection RNA-sequencing data are obtained from the GEO database (accession id GSE157490). The Spycone package is available as a PyPI package. The source code of Spycone is available under the GPLv3 license at https://github.com/yollct/spycone and the documentation at https://spycone.readthedocs.io/en/latest/. The code used to produce the result shown in this manuscript is compiled into the Google colab notebook (https://colab.research.google.com/drive/13CjfzZizPlmxzsT-zm6zEgfdFce1fzSC?usp=sharing). This workflow is documented in the AIMe registry https://aime.report/DXKacH [57].

## Ethics approval and consent to participate

Not applicable.

## Author’s contributions

TK, OT, JB, and ML conceived the project. CTL performed the initial experiments, developed the method, and implemented the software. OT, TK and ML supervised the project, provided critical feedback, and helped shape the research. CTL wrote the initial draft of the manuscript. ZL and AF contributed to the discussions and method development. All authors contributed to writing the final manuscript and approved the final version.

## Acknowledgements

Not applicable.

