## Supplementary figures for "Systematic analysis of alternative splicing in time course data using Spycone"

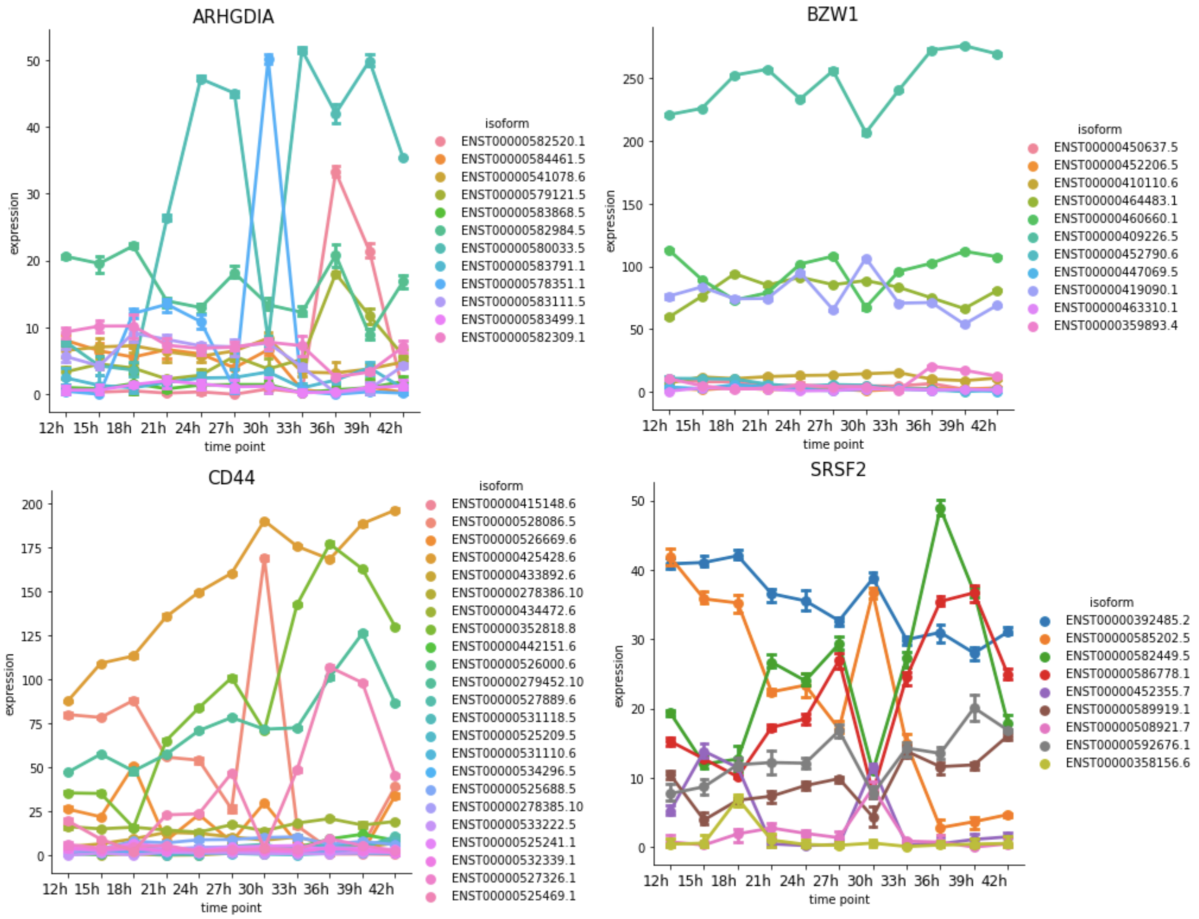


S1. Real life IS examples that are challenging to detect. In the case of gene ARHGDIA, at 30h a lowly expressed isoform (light blue) is up-regulated while the highest abundance isoform. In the case of CD44, a lowly expressed isoform (pink) is up-regulated at 30h, however, there is no other isoform down-regulated with the similar magnitude as in ARHGDIA. Similar case in SRSF2, with an isoform (orange) up-regulated at 30h, multiple isoforms down-regulated at the same time point. This case could be an example of multiple isoform switching. In BZW1 gene, two isoforms seem to switch at 30h and 33h, however, in the earlier time points, the two isoforms are expressing parallelly.

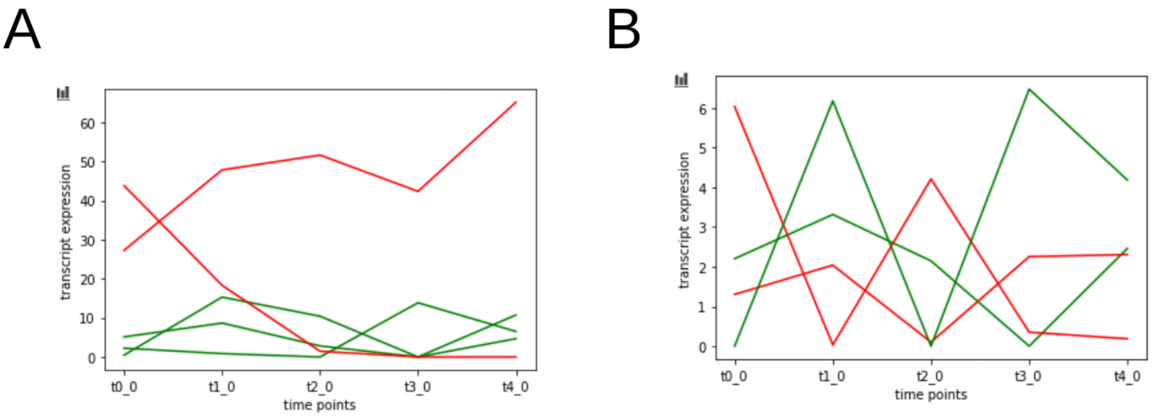


S2. Model 1 and Model 2 examples from simulated data. Model 1 allows switching only with the highest abundance isoform. Model 2 allows isoforms with higher than 0.3 abundance to switch.

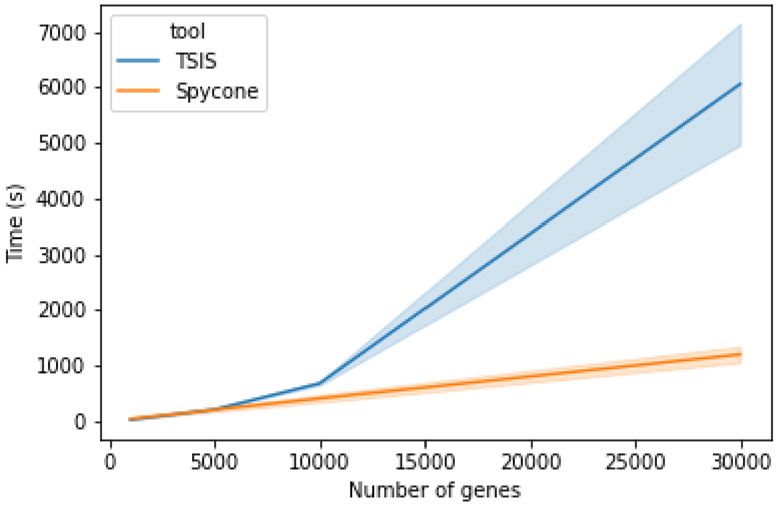


S3. Computation time for Spycone and TSIS (seconds) on simulated datasets ranging from 1000 to 30000 genes with 3 to 4 replicates. Tested on a laptop device with Intel CoreTM i7-10510U CPU @ 1.8GHz x 8 cores, 16GB RAM, 512GB disk capacity.

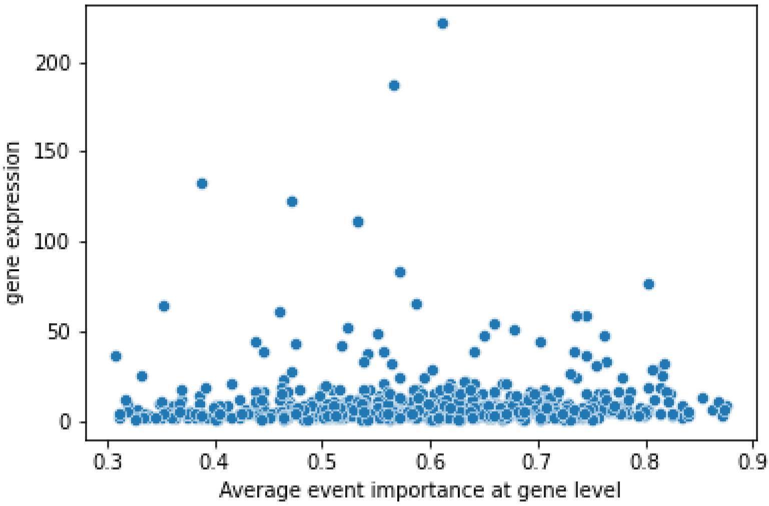


S4. Relationship between event importance and overall gene expression from Spycone results.

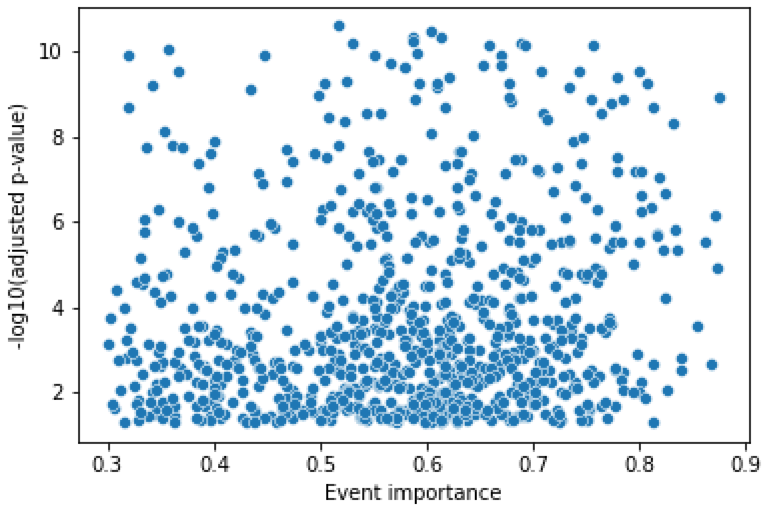


S5. Relationship between event importance and adjusted p-value from Spycone results.

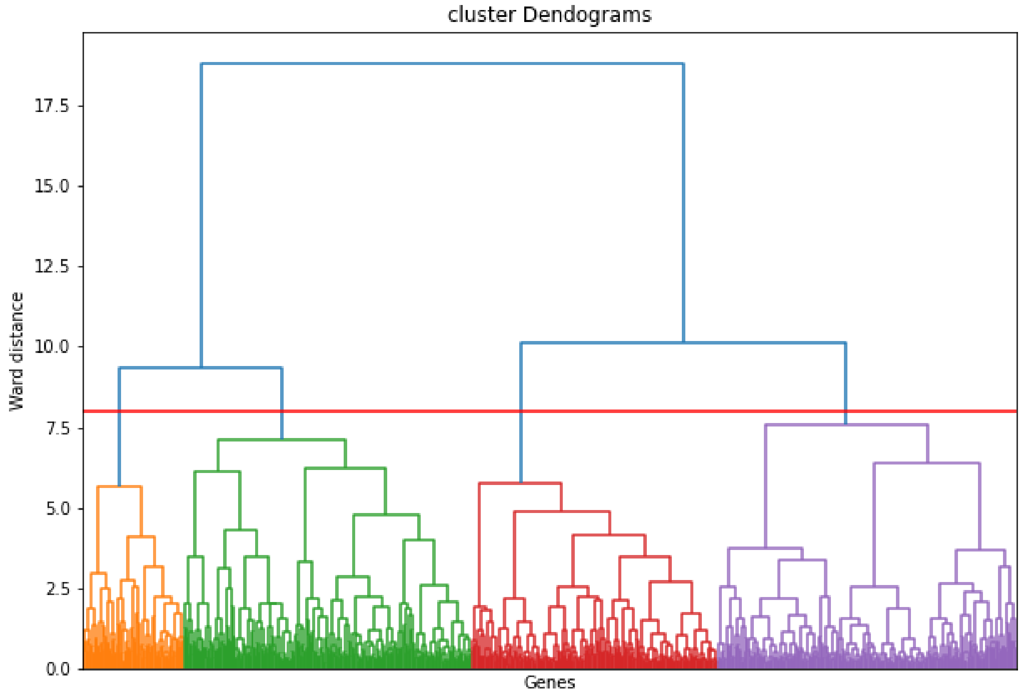


S6. Cluster dendrogram of hierarchical clustering SARS-Cov-2 dataset. Each cluster is colored with different colors under the ward distance threshold at 8.

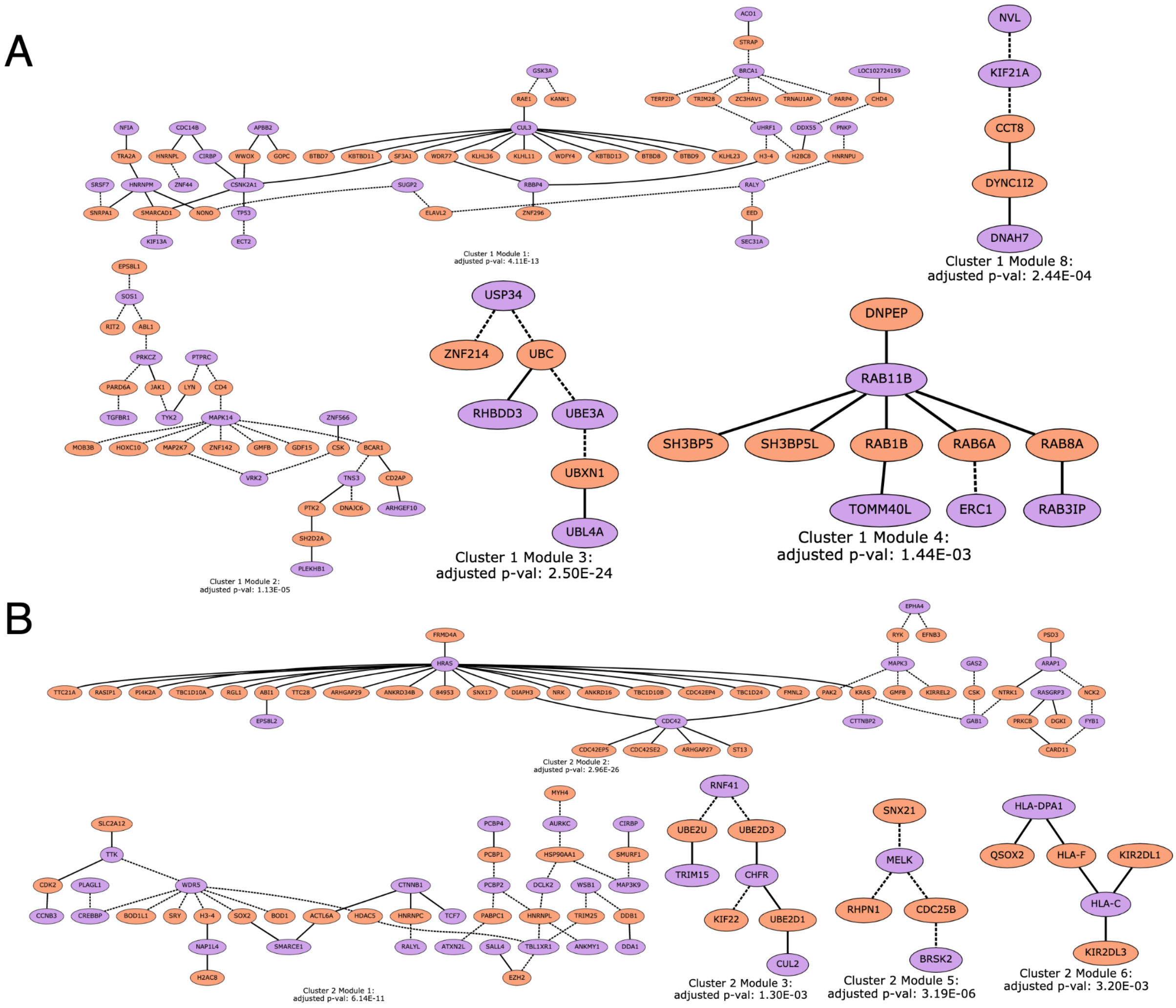


S7. Spycone results in modules of PPI network and their gene set enrichment results. Active network modules are identified using DOMINO. Each node represents a domain of a gene. Purple nodes are the isoform switched genes and orange nodes are non-IS genes from the PPI. Dashed edges are the affected interactions between the genes due to the lost/gained of domains during the IS events.

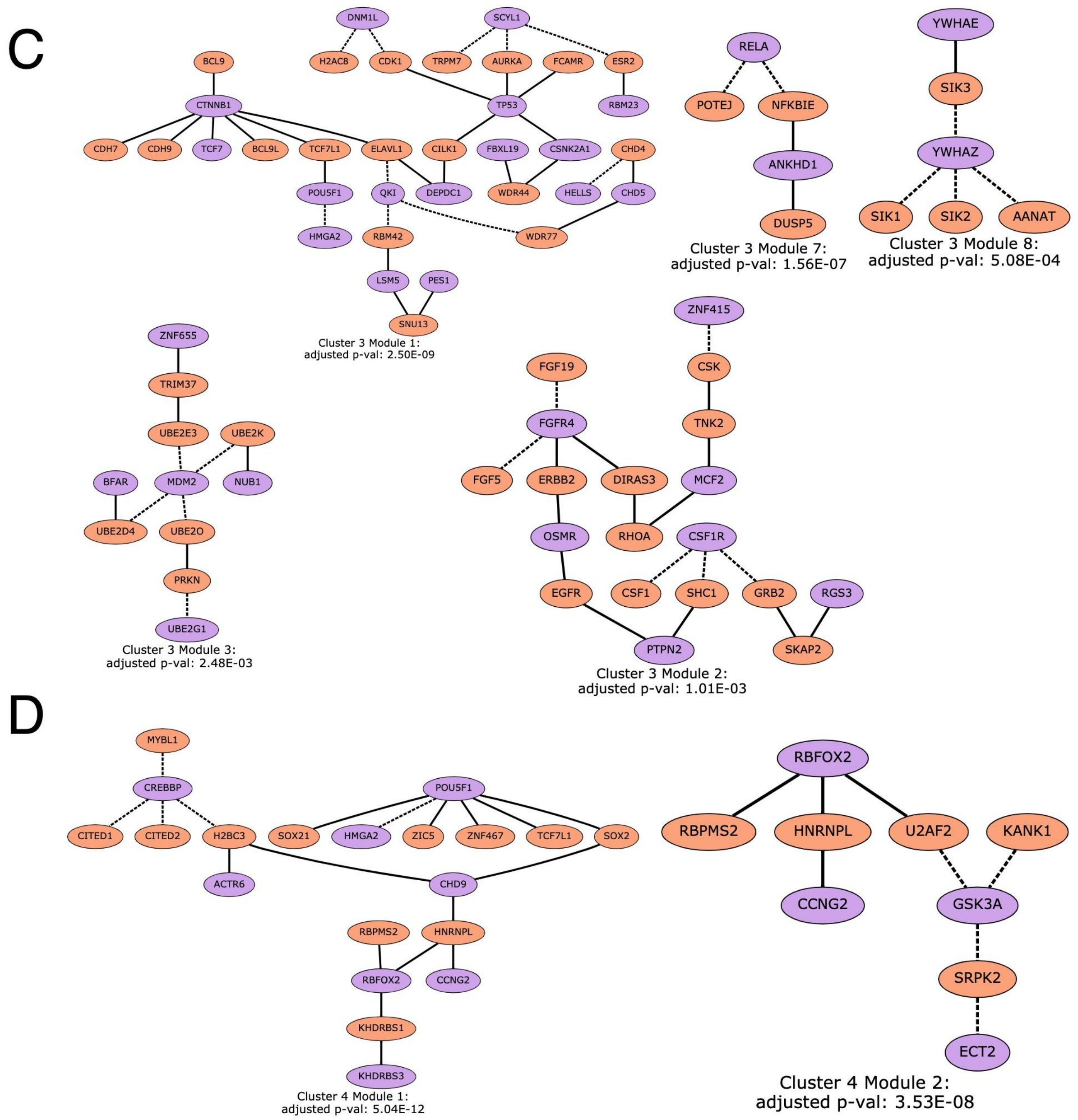


S7 (Cont). Spycone results in modules of PPI network and their gene set enrichment results. Active network modules are identified using DOMINO. Each node represents a domain of a gene. Purple nodes are the isoform switched genes and orange nodes are non-IS genes from the PPI. Dashed edges are the affected interactions between the genes due to the lost/gained of domains during the IS events.

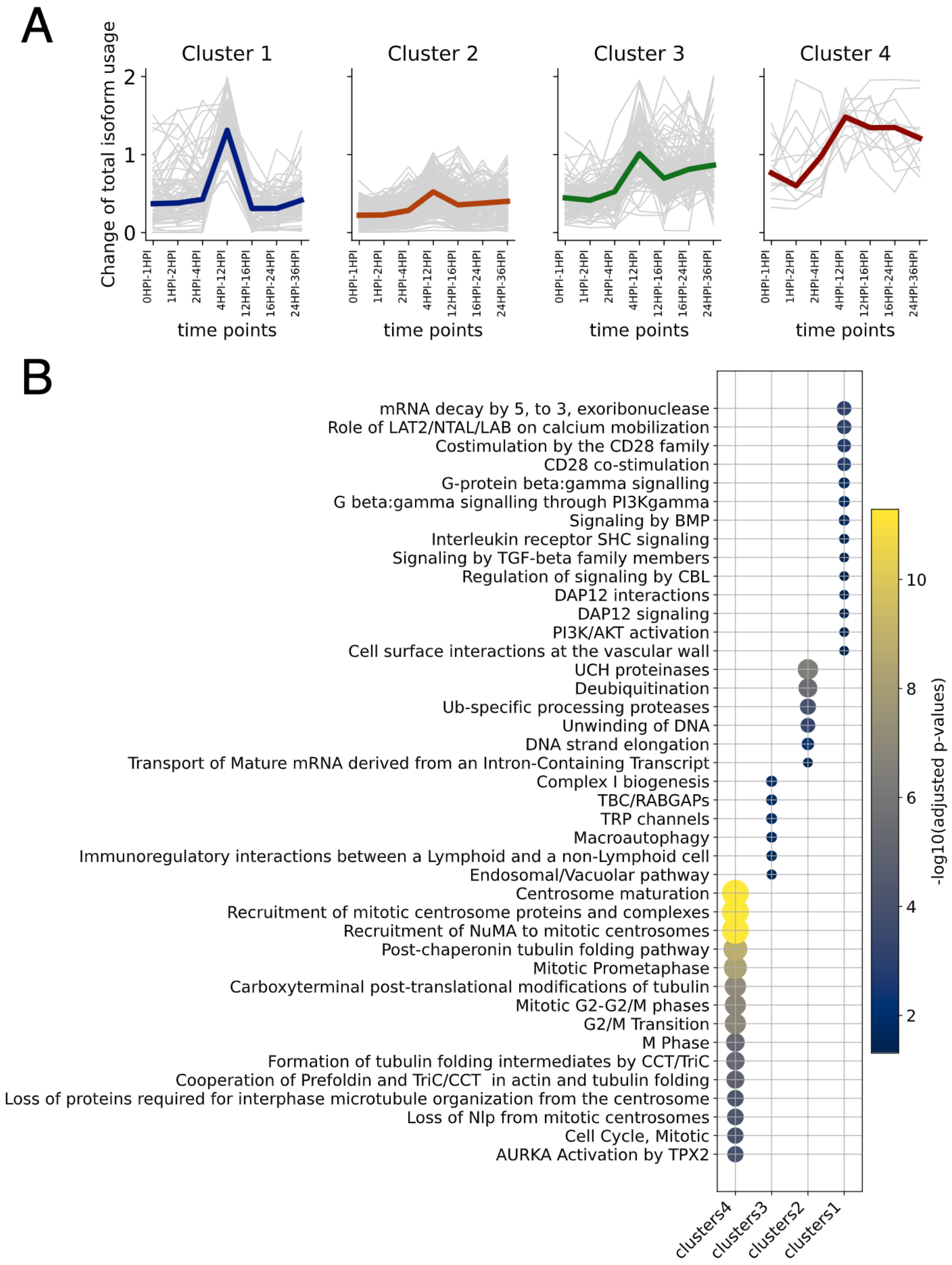


S8. Clustering and NEASE results from TSIS identified IS genes.
