## Supplementary information - use case 2 for "Systematic analysis of alternative splicing in time course data using Spycone"

### Application to human cancer cell line data

Here we describe a Spycone analysis of a human HD-MY-Z tumor cell line RNA-seq dataset with 11 time points and no replicates [1].

We used Trimmomatic for adapter trimming and Salmon for isoform quantification with human genome version 38, and an Ensembl genome annotation version 103. We kept isoforms with TPM > 1 across all time points and used 47577 isoforms to detect IS events.

Only the results of Spycone are shown since TSIS required at least 3 replicates. Spycone reported 345 pairs of isoforms: 152 of them affect at least 1 domain. The event importance distribution is shown in Fig. 1A.

We calculated the changes in total isoform usage across consecutive time points, applied ward linkage hierarchical clustering, and obtained 4 clusters (Table S6, Figure 1B, C). Then we performed functional enrichment with NEASE using the Reactome (Fig. 1D) and KEGG pathway databases (Table S7) as described above. Cluster 1 has a higher change of total isoform usage in the later stage, where cell cycle (G2/M phase) progression is affected. Cluster 2 has a similar pattern as cluster 1 with lower change of total isoform usage. It is enriched in post-translational modification and early elongation complex. Cluster 3 has higher change of total isoform usage across time points, it is enriched in splicing, chromatin remodeling related pathways, and rRNA expression. Cluster 4 has less change of total isoform usage across time points, where intra-cellular trafficking (Golgi apparatus related) and siRNA and miRNA biogenesis are enriched.

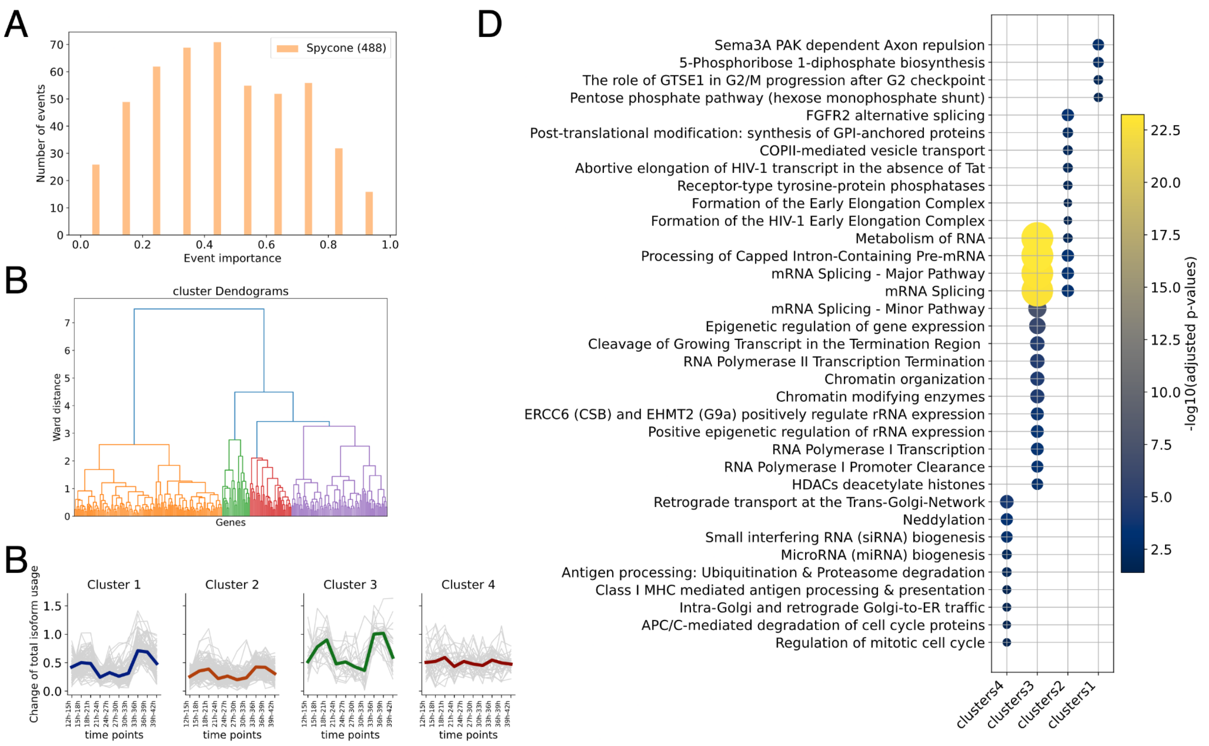


Figure 1. IS detection results from Spycone A) The distribution of the detected events based on the event importance metric. Number of events is indicated inside the brackets in the legend. B) Cluster dendrogram of hierarchical clustering. Each cluster is colored with different colors under the ward distance threshold at 3.3. C) Cluster prototypes (colored line) and all objects (gray lines) show the pattern of the change of total isoform usage across time points. D) NEASE enrichment result of the IS events in each cluster.

### Gene modules with transcription factors affected by IS events

We detected active modules as described in the main text. Examples of the resulting networks with significant bonferroni adjusted p-value are shown in Fig. 2.

We found 2 modules in cluster 1 (Fig. 2A). Module 1 contains RNA-binding proteins like SRSF5, SF3B4, SRSF2, TRA2A, TRA2B, which are performing mRNA splicing and processing. Module 2 consists of genes related to ubiquitination.  We found 3 modules in cluster 2 (Fig. 2B). In module 1, many genes are transcription factors such as TCF19, CREBBP, CITED2, DDX17 and TAF15.  Interestingly, SRPK2 has affected interaction with YPEL3, a gene related to cell senescence and PHLDB1 that regulates epithelial to mesenchymal transition. These processes induce cell cycle arrest in which SRPK2 is found to be regulating in non-small cell lung cancer [2]. In module 3, PTPRF, a tyrosine phosphatase has affected interaction with RhoA associated kinase ROCK2 and Interleukin IL13RA2. The interplay of interleukin and ROCK signaling might contribute to cancer growth as suggested in previous studies [3,4]. There is 1 module found in cluster 3 (Fig. 2C). RAB7A regulates vesicle trafficking, and it is activated by MON1A-MON1B-CCTZ guanine exchange factor (GEF) complex. RAB7A is part of Ras oncogene family, and its knockdown suppresses tumor growth in breast cancer [5]. There are 2 modules found in cluster 4 (Fig. 2D). Module 1 contains mostly RNA binding proteins: splicing factors (MAGOH, TRA2B, RBFOX2, SNRNP70, HNRNPL), RNA processing (TUT4, MATR3, RBM3, RBMS1), as well as transcription factors (YAP1, MSI2) and chromatin remodelers (RECQL4, SMYD5). TRIM25 functions as a ubiquitin E3 ligase, which is associated with breast cancer [6,7]. Module 2 is found with ribosome-related proteins with RNA binding activity (NOP2, NIP2). NOP2 acts as a target of lncRNAs which leads to hepatocellular carcinoma cell proliferation and prostate cancer proliferation [8,9].

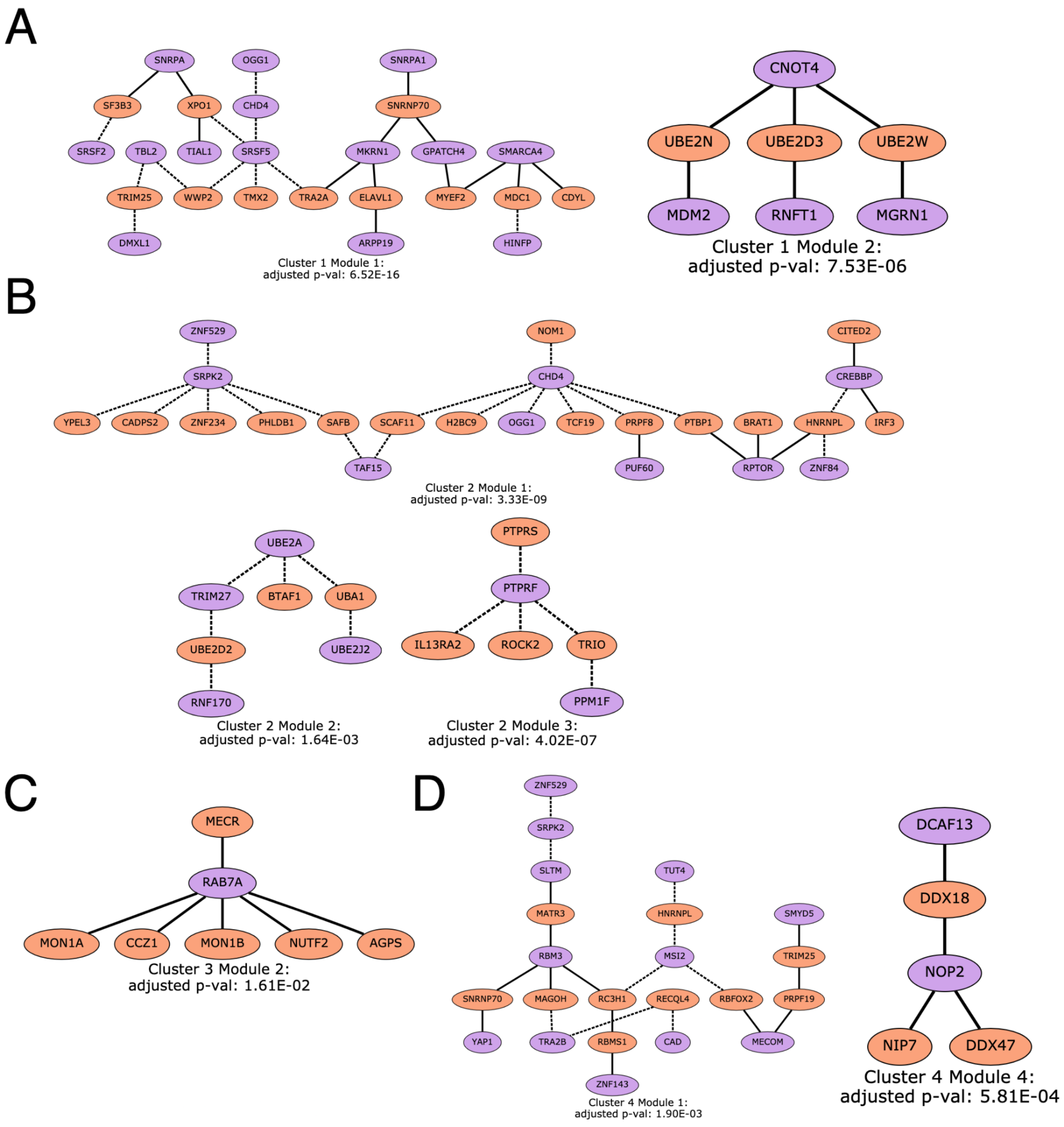


Figure 2. Spycone results in modules of PPI network and their gene set enrichment results. Active network modules are identified using DOMINO. Each node represents a domain of a gene. Purple

nodes are the isoform switched genes and orange nodes are non-IS genes from the PPI. Dashed edges are the affected interactions between the genes due to the lost/gained of domains during the IS events.

**Splicing factor analysis**

As described in the main text, we performed co-expression analysis and motif search using PSSM to determine potential splicing factors that regulate IS events during tumor development. We found six significant RNA-binding proteins that are co-expressed with at least two isoforms of the same gene in the cluster (adjusted p-value < 0.05) (Table S10).
To investigate the binding potential of the alternative spliced exons during IS events within a cluster, we perform RNA-binding proteins' motifs enrichment analysis (Table S11). The IGF2BP1, CSTF2_0, CSTF2_3, DDX3X_7 and PPIG_1 motifs show higher log-odd scores at the 5' end of the lost/gain exons than of the unregulated exons (one-sided Mann-Whitney U test p-value < 0.05) (Fig. 3A). CSTF2_0, CSTF2_3 and DDX3X_7 motifs show higher log-odd scores at the 3' end in cluster 1 (Fig. 3B). IGF2BP1 is identified as a mRNA stabilizer to promote cancer cell progression [10]. CSTF2 is found inducing the shortening of RAC1 isoform at the 3'UTR in human urothelial carcinoma [11] and various genes in a cancer cell line [12]. DDX3X directly interacts and regulates the splicing of KLF4 in a cancer cell line [13].

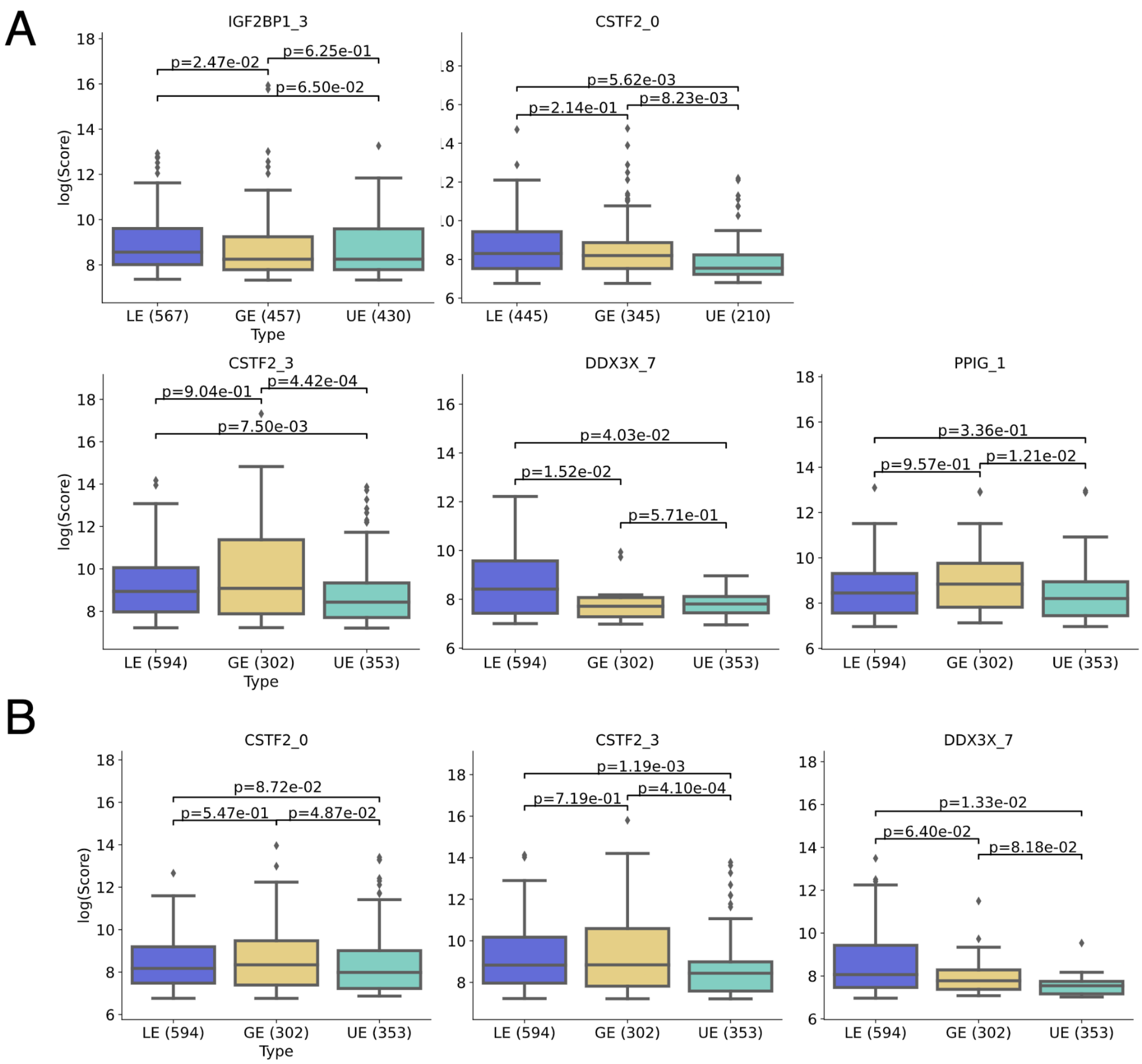


Figure 3. A) The motif logos of the NCBP2_11 motif from WebLogo. B) Boxplots showing the PSSM scores difference between lost/gained exons and background exons at exons 5' boundaries with a score threshold of 10 in logarithmic scale. (one-sided Mann Whitney U test p-value < 0.05). C) PSSM scores difference between lost/gained exons and background exons at exons 3' boundaries with a score threshold of 10 in logarithmic scale. (one-sided Mann Whitney U test p-value < 0.05).

2. Li X, Yang S, Zhang M, Xie S, Xie Z. Downregulation of SRPK2 promotes cell cycle arrest

though E2F1 in non-small cell lung cancer. Eur J Histochem [Internet]. 2019;63. Available from: http://dx.doi.org/10.4081/ejh.2019.3067

13. Cannizzaro E, Bannister AJ, Han N, Alendar A, Kouzarides T. DDX3X RNA helicase

affects breast cancer cell cycle progression by regulating expression of KLF4 [Internet]. FEBS Letters. 2018. p. 2308–22. Available from: http://dx.doi.org/10.1002/1873-3468.13106
